# Contextual Evaluation of MicroRNA Sequencing Data Harmonization: Performance in Sample Clustering

**DOI:** 10.64898/2026.08.09.743718

**Authors:** Jian Zou, Yannick Düren, Xinyi Wang, Ying Xiang, Yunhui Qi, Miao Wang, Yilin Wu, Samuel Singer, Li-Xuan Qin

## Abstract

Reliable translation of microRNA sequencing data depends on effective harmonization to mitigate artifacts from variable experimental handling. Although many harmonization methods exist, prior evaluations have focused mainly on differential expression analysis, leaving the impact on subgroup discovery understudied. We present a framework for evaluating harmonization in the context of sample clustering, which integrates AI-augmented datasets, statistical evaluation pipelines, and accessible software tools, enabling systematic comparisons across diverse signal-to-artifact ratios and cluster composition settings. Using this framework, we show that harmonization can, often partially, restore clustering accuracy lost to artifacts, especially at moderate signal-to-artifact ratios, with the level of gains depending on the specific harmonization method, the paired clustering technique, and the cluster composition setting. We further confirm these findings by analyzing reconstructed cohorts from The Cancer Genome Atlas breast cancer microRNA sequencing data. Collectively, the results underscore the need for tailored harmonization to support reliable subgroup discovery and highlight the broader importance of context-specific workflows in translational genomics.

## Introduction

MicroRNAs (miRNAs) are small, non-coding RNAs that play crucial roles in regulating gene expression^1,2^. Their involvement in key cellular processes, such as cell differentiation, apoptosis, and carcinogenesis, heralds their potential as powerful biomarkers for disease diagnosis and prognosis^3,4^. Similar to RNA sequencing, miRNA sequencing data analysis faces significant challenges from unwanted variations introduced during experimental handling^5,6^. If left unaddressed, these artifacts can lead to false discoveries and obscure true biological signals, prompting the need for effective harmonization that maximally reduces artifacts while best preserving signals to ensure reliable data-to-knowledge translation^7,8^.

Many existing methods for data harmonization (including normalization and batch-effect correction) were developed for RNA sequencing and later applied to miRNA sequencing, yet their performance often differs, as we recently demonstrated^8,9^. These differences partly reflect the unique features of miRNA data, like strong tissue specificity and low molecular complexity (i.e., a few dominantly abundant markers)^10–13^. Beyond data type, harmonization effectiveness also depends on the downstream analysis, as each task defines “biological signal” differently and can be influenced by harmonization in distinct ways, a pattern already noted for miRNA microarrays^14–19^. We hypothesize that this analysis-context dependence extends to miRNA sequencing. Prior work on its harmonization performance, including ours, has focused largely on differential expression analysis^5,7–9^. Here, together with a complementary back-to-back study, we evaluate performance in two additional analysis contexts: sample clustering and classification.

Our investigation leveraged a unique pair of miRNA sequencing datasets for the same set of tumor samples collected at Memorial Sloan Kettering Cancer Center (MSK)^8^. The first dataset was collected under a carefully controlled study design, with not only uniform handling of all samples by the same technician in a single experiment run but also balanced assignment of samples to multiplexed sequencing libraries via blocking and randomization, aiming to minimize artifacts and prevent confounding. In contrast, the second dataset was collected following typical practice (that is, by multiple technicians over an extended period and in the order of sample collection), exhibiting evident artifacts. To enhance generalizability, we employed deep generative models to increase both the sample size and the number of paired datasets^20^, and then simulated scenarios over a range of artifact magnitudes under various signal strengths (Fig. 1).

**Fig. 1.**
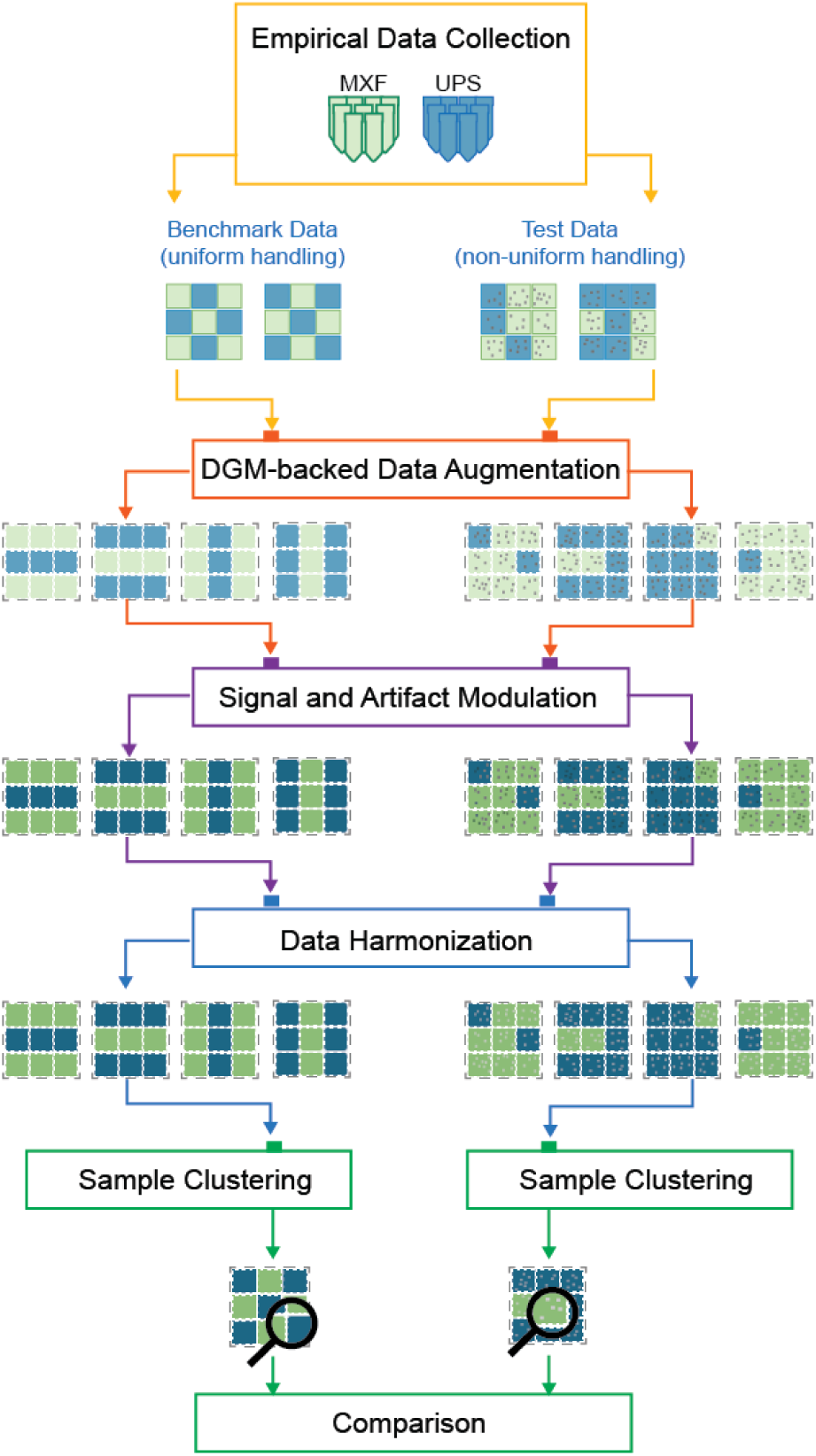
Study overview. Paired microRNA sequencing datasets were collected for the same set of 27 Myxofibrosarcoma (MXF) samples and 27 Undifferentiated Pleomorphic Sarcoma (UPS) tumor samples. The benchmark dataset was generated with uniform handling and balanced library assignment, whereas the test dataset was processed by multiple technicians over several years in order of sample collection. This dataset pair was then input to a deep-generative-model-powered data augmentation algorithm to create additional paired dataset of varying sample sizes. Their biological signal strength and artifact magnitude were subsequently modulated, and the resulting paired datasets were used to evaluate multiple harmonization methods in combination with clustering techniques.

Our findings show that harmonization performance varies across signal-to-artifact ratios and clustering techniques, and that no single method dominates overall. For instance, quantile normalization (QN) produced high accuracy for most clustering techniques when the ratio was relatively high, and it maintained solid performance at moderate ratios especially with hierarchical clustering using selected distance metrics. Notably, however, QN was among the worst performers in differential expression analyses, particularly at strong or modest ratios^8^. These contrasts underscore the need to align harmonization with the specific analytical context and even the chosen analysis approach for a given signal-to-artifact setting. Applied to The Cancer Genome Atlas (TCGA) breast cancer data^21^, harmonization again substantially affected clustering performance, with the magnitude of this effect depending on both the harmonization method and the analysis approach. These results highlight the pronounced interplay between data harmonization and downstream analysis and the potential to improve analytical outcome through tailored harmonization.

Given the important role of sample clustering in miRNA sequencing data analysis for uncovering novel subgroups and refining disease diagnosis, effective harmonization is vital for harnessing biological insights and advancing clinical translation. To enable rigorous, reproducible, and extensible evaluation, we developed the *PRECISION.seq.augmented* R package, which implements our framework using realistically distributed, robustly benchmarked, and flexibly configured data. This open-source resource allows users to reproduce our analyses and extend them to additional harmonization methods and clustering techniques.

## Methods

### Collection of empirical data

Myxofibrosarcoma (MXF) and Undifferentiated Pleomorphic Sarcoma (UPS; previously known as Pleomorphic Malignant Fibrous Histiocytoma, PMFH) are two aggressive subtypes of soft tissue sarcoma^22^. We analyzed 54 newly diagnosed, untreated primary tumor samples – 27 MXF and 27 UPS – collected at MSK between 2000 and 2012, following prior protocols^8,9^. Each sample was sequenced twice, yielding two datasets: (1) the benchmark dataset, processed with uniform handling and balanced library assignment, and (2) the test dataset, collected by multiple technicians over several years as specimens became available. Due to the extended collection period, the test dataset did not retain an explicit batch variable.

### Augmentation of empirical data using deep generative models

We adapted the SyNG-BTS^20^ algorithm to expand both the sample size and the number of paired datasets. Recently developed to augment bulk-tissue transcriptomic sequencing (log2 count) data from small pilot studies, SyNG-BTS showed robust performance across multiple assessment datasets and metrics (marker-level summaries, inter-marker correlations, and sample-level similarities), under varied generative models, hyperparameter settings, pilot study sizes, and preprocessing schemes^20^. Here, the paired 54-sample datasets served as input pilot data, with the two data vectors for each sample concatenated into a single super vector to preserve the benchmark-test pairing. We used a Masked Autoregressive Flow model with the following hypermeters: batch fraction of 0.1, learning rate of 0.0005, and 160 training epochs with early stopping (i.e., training halted if validation loss failed to improve for 20 consecutive epochs) to prevent overfitting and avoid unnecessary computation. To confirm congruence between the empirical and augmented data, we randomly selected 30 augmented datasets and graphically compared them with the empirical dataset at the marker level using scatter plots of summary statistics (such as group-specific means and standard deviations) and at the sample level using Uniform Manifold Approximation and Projection (UMAP)^23^. Additionally, for each augmented dataset, we summarized concordance with the empirical dataset at multiple levels: (i) inter-sample, defined as mixing of generated and real samples measured by the complementary Adjusted Rand Index (cARI); (ii) inter-marker, defined as similarity of inter-marker correlations among miRNAs within the same polycistronic clusters (PCCs), quantified by concordance correlation coefficients (CCCs); and (iii) marker-level, measured by similarity in between-group mean differences and by similarity in benchmark-versus-test correlations, each also quantified by CCCs.

### Modulation of signal strength and artifact magnitude in paired augmented datasets

For a given augmented dataset pair, let X*_MXF_* and X*_UPS_* represent the benchmark data matrices for MXF and UPS samples, respectively, and let *Y_MXF_* and *Y_UPS_* represent the corresponding test data matrices. Denote the combined matrices as X = [X*_MXF_* X*_UPS_*] and *Y* = [*Y_MXF_ Y_UPS_*] . Let *X_MXF_* and *X_UPS_* denote the marker-specific means in the benchmark data for MXF and UPS, respectively.

Biological signals were defined as the differences between MXF and UPS samples in the benchmark dataset, denoted by *̄X_MXF_* – *̄X_UPS_*. To amplify these signals, [*̄X_MXF_* – *̄X_UPS_*] ⁢ *c* was added, for each marker, to the samples in the group with the greater mean in the benchmark dataset. The same operation was applied to the paired test dataset to ensure consistent signal levels.

Under the assumption of additivity on a log-transformed scale^14, 24–26^, we defined handling artifacts in a test dataset as its differences from the paired benchmark dataset, denoted as *Y* − X. To modulate the magnitude of artifacts, (*Y* − X) × *d* was added to *X* to form the modulated test dataset.

We considered five levels of biological signal strength – very weak (*c* = 0.2), weak (*c* = 0.6), moderate (*c* = 1.0), strong (*c* = 1.5), and very strong (*c* = 2.0) – and three levels of technical artifact magnitude – weak (*d* = 1.2), moderate (*d* = 1.5), and strong (*d* = 2.0). Of note, the strong and very strong signals levels were omitted for clustering analysis of benchmark datasets, as clusters were already nearly perfectly separated at the moderate level, whereas the very weak signal level was excluded when clustering test datasets, as clustering at the weak signal level was already near random in the presence of weak artifacts. For each simulation scenario, 300 datasets were generated, each consisting of 100 MXF and 100 UPS samples, reflecting the typical sample sizes in TCGA^27^. To assess the impact of cluster size imbalance, we generated additional datasets with skewed class distributions, such as 20 MXF samples and 180 UPS samples.

### Data harmonization

To mitigate unwanted variations, we applied two types of harmonization methods to the modulated data: scaling-based and regression-based. The former included Total Count (TC; *edgeR::cpm*)^7^, Upper Quartile (UQ)^5^, Median (Med)^7^, Trimmed Mean of M-values (TMM; *edgeR::calcNormFactors(“TMM”)*)^28^, DESeq (*DESeq2::estimateSizeFactors*)^29^, and PoissonSeq (*PoissonSeq::PS.Est.Depth*)^30^. The latter included Quantile Normalization (QN; *preprocessCore::normalize.quantiles*)^31^ and Remove Unwanted Variation (RUV; *RUVSeq*) with its three variants – RUVg, RUVr, and RUVs^32^. RUV estimates latent batch effects via factor analysis and adjusts for these factors within a linear-regression framework; we used one latent factor estimated from the 85% randomly selected markers as a default and explored additional settings in sensitivity analyses. For comparison, we also included no harmonization as a baseline.

### Sample clustering and evaluation

To re-discover sample subgroups (i.e., tumor subtypes) in the data, we applied five clustering techniques: K-means^33^, Partitioning Around Medoids (PAM)^34^, Self-Organizing Maps (SOM)^35^, Multivariate Normal Mixture models (MNM)^36^, and Hierarchical Clustering (HC)^37^. Both PAM and HC were implemented with three different distance metrics – Euclidean distance, Pearson correlation, and Spearman correlation – denoted as -E, -P, and -S, respectively. Among these techniques, K-means, PAM, and SOM represent algorithm-based partitional clustering, whereas MNM is model-based partitional and HC is algorithm-based hierarchical. These techniques were implemented in R (version 4.4.1) using the following functions: *kmeans* for K-means, *pam::cluster* (v2.1.6) for PAM, *som::som* (v0.3-5.2) for SOM, *Mclust::mclust* (v6.1.1) for MNM^38^, and *hclust* for HC. The resulting clusters were compared with tumor subtypes using the Adjusted Rand Index (ARI)^39,40^. The ARIs for the 300 datasets in each simulation scenario were averaged and visualized as heatmap-style tables; corresponding 95% confidence intervals are provided in the Supplementary Materials.

### Analysis of reconstructed TCGA cohorts

To further illustrate the benefit of tailored harmonization in real-world data, we reanalyzed the miRNA sequencing data from the TCGA breast cancer (BRCA) study, whose default harmonization is TC scaling^41^. We retrieved read-count files for 994 primary tumor samples using the *TCGAbiolinks* R package^42^, and downloaded breast cancer subtype labels (Luminal A, Luminal B, Basal-like, and HER2) from Ellrott *et al*^43^. We extracted data-collection dates using the *MBatch* package and categorized them into early (from June 2010 to early May 2011), middle (from late May to early August 2011), and late batches (from late August 2011 to May 2014), based on the observed temporal distribution of sequencing depth. The middle batch was treated as a transition period and excluded from cohort reconstruction.

Given the available sample sizes and the degree of miRNA expression differences between subtypes, we focused on Luminal A and Basal subtypes within the early and late batches, totaling 641 samples. From these, we constructed four cohorts to represent distinct artifact scenarios: (I) a weak artifact scenario, with all samples drawn from the late batch; (II) a strong, balanced artifact scenario, with equal numbers of early- and late-batch samples for each subtype; (III) a strong, completely confounded scenario, with all Luminal A samples from the early batch and all Basal samples from the late batch; and (IV) a parallel confounded scenario with the batch assignment switched between subtypes (i.e., all Luminal A samples from the late batch and all Basal samples from the early batch). Exact sample sizes by subtype and batch for each reconstructed cohort are provided in Table S1. For each scenario, we generated 10 datasets through resampling (with replacement). Each dataset was then harmonized and clustered using the same methods as in our simulation study.

## Results

### Augmented data closely mimic empirical data

We first evaluated the augmented data graphically at both the sample and marker levels (Fig. 2A-2D). At the sample level, generated samples co-located with real samples on UMAP, indicating strong concordance. At the marker level, marker-specific means and standard deviations of the generated samples (with each sample type) aligned closely with those of the real samples, preserving the original mean-variance relationship without inflating or attenuating dispersion.

**Fig. 2.**
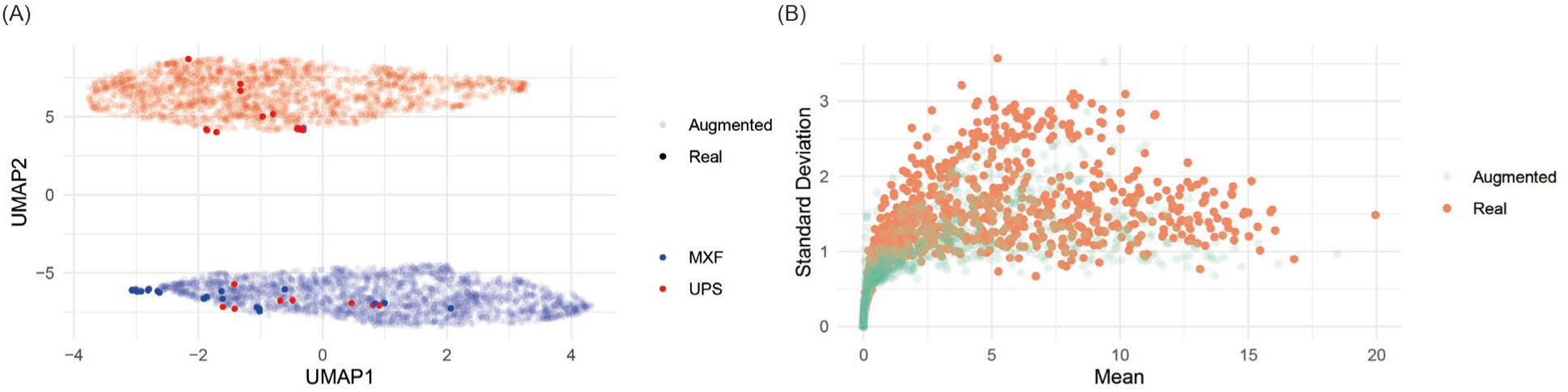
Quality assessment of augmented data. (A) Marker-level mean-versus-standard deviation plot indicates comparable distributional properties between empirical and augmented datasets for both MXF and UPS groups in the benchmark dataset. (B) UMAP of sample profiles for the benchmark dataset shows generated samples co-locating with real samples for both MXF and UPS groups. (C) Marker-level mean-versus-standard deviation plot for the test dataset likewise indicates comparable distributional properties between empirical and augmented data. (D) UMAP of sample profiles for the test dataset likewise shows generated samples co-locating with real samples for both MXF and UPS groups. (E) Boxplots of cARI values across augmented datasets (each compared with the empirical dataset) show good mixing of generated and real samples. (F–H) Boxplots of CCCs across augmented datasets (each compared with the empirical dataset) show good preservation of (F) inter-marker correlation structures among miRNAs within polycistronic clusters, (G) marker-specific between-group mean differences, and (H) marker-specific correlations between each benchmark dataset and its paired test dataset.

We further quantified concordance between each augmented dataset and the empirical dataset using four numerical metrics and displayed them as box plots (Fig. 2E-2H). At the inter-sample level, generated and real samples mixed well, as reflected by high cARI values (Inter-quartile range [IQR]: 0.967 to 0.996 for benchmark MXF, 0.911 to 0.987 for benchmark UPS, 0.983 to 0.996 for test MXF, and 0.978 to 0.997 for test UPS). At the inter-marker level, miRNAs within PCCs showed similar correlation structures in augmented and empirical data, reflected by strong CCCs (IQR: 0.725 to 0.756 for benchmark and 0.755 to 0.775 for test). At the marker level, between-group mean differences were largely preserved, again indicated by strong CCCs (IQR: 0.816 to 0.886 for benchmark and 0.795 to 0.874 for test). Finally, we introduced an additional marker-level metric to assess correlation between each benchmark dataset and its paired test dataset and found good preservation in generated samples relative to real samples (IQR: 0.701 to 0.723).

### Harmonization may slightly improve or inadvertently worsen clustering in the absence of artifacts

To assess the effect of harmonization on clustering in the absence of artifacts, we analyzed the augmented benchmark datasets before and after harmonization across three levels of biological signal strength (Fig. 3; Table S2).

**Fig. 3.**
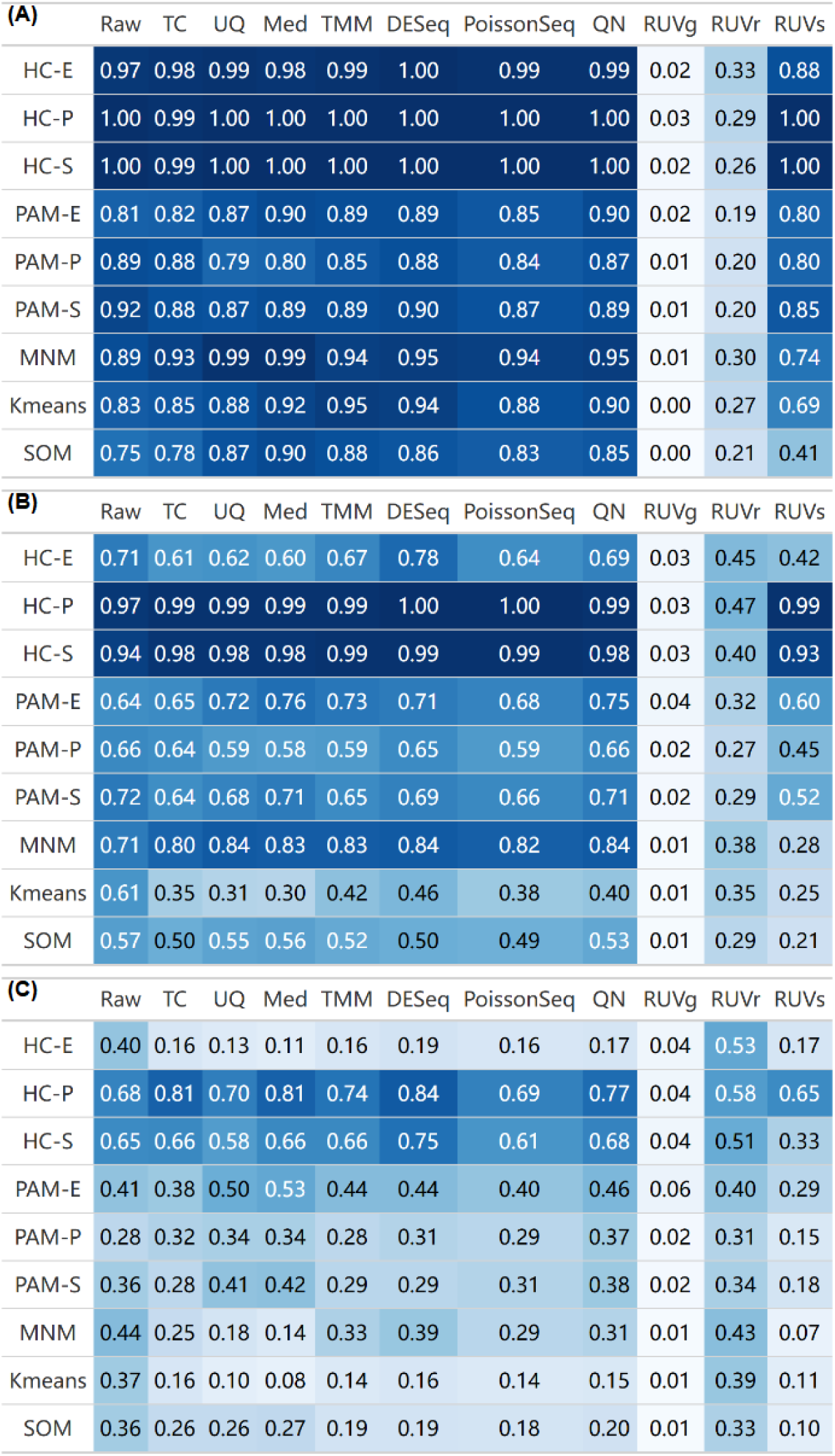
Evaluation of harmonization performance for clustering using augmented data without artifacts. Heatmaps show mean Adjusted Rand Index (ARI) values for different harmonization methods combined with various clustering techniques across three levels of biological signal strength: (A) c = 1, (B) c = 0.6, and (C) c = 0.2. Higher ARI values (darker blue) indicate better clustering accuracy.

#### Moderate signals

When the two sample groups were well-separated (*c* = 1.0), hierarchical clustering achieved near-perfect separation without harmonization, regardless of the distance metric. Specifically, HC- P and HC-S each reached an ARI of 1.00, while HC-E close behind at 0.97. Scaling-based harmonization did not affect the accuracy of HC-P and HC-S, and it improved HC-E toward 1.00.

Prior to harmonization, PAM-P and PAM-S attained ARIs of 0.89 and 0.92. After scaling harmonization, both declined slightly; for example, UQ lowered them to 0.79 and 0.87, respectively. By contrast, PAM-E, K-means, MNM, and SOM started at 0.75–0.89 before harmonization and benefited from scaling, with gains up to 0.15; for instance, TMM increased SOM from 0.75 to 0.88.

Among regression-based methods, QN performed on par with the better scaling methods. However, the RUV variants reduced clustering performance: RUVs caused small-to-moderate declines, while RUVr and RUVg led to substantial deterioration, with ARIs falling to ∼0.25 and <0.03, respectively. For illustration, Fig. S1 shows the effect of RUVg on the clustering structure, compared with TMM, in an example dataset.

#### Weak signals

When the sample groups were less well-separated (*c* = 0.6), clustering performance declined across all harmonization methods (except RUVr and RUVg) and clustering techniques, as expected. Relative to the moderate-signal scenario, ARIs dropped the least for MNM and the most for K-means and SOM.

Without harmonization, HC-P (0.97) and HC-S (0.94) again led in ARI, while the other techniques ranged from 0.57 to 0.72. With scaling-based methods, HC-P and HC-S remained top performers, approaching perfect ARIs, and MNM emerged as the next best, rising from 0.71 to 0.80–0.84.

Several scaling methods notably improved mid-tier techniques. For instance, MNM’s ARI rose from 0.71 to 0.84 under DESeq and UQ. Conversely, low-tier techniques such as K-means and SOM deteriorated markedly, with ARIs dropping from 0.61 and 0.57 to approximately 0.30–0.50 after scaling.

Among regression-based methods, QN showed mixed effects, enhancing clustering for some techniques (e.g., MNM and PAM-E) but worsening it for others (e.g., K-means). RUV variants continued to degrade ARIs across clustering techniques, with the exception of RUVs used with HC-P (which slightly improved from 0,97 to 0.99). Notably, RUVg essentially erased the clustering structure.

#### Very weak signals

When the two groups overlapped substantially (*c* = 0.2), ARIs fell sharply regardless of harmonization. Without harmonization, even the top-performers, HC-P and HC-S, dropped to 0.68 and 0.65, respectively, while most other techniques hovered around 0.40.

With scaling or QN, HC-P and HC-S improved (up to 0.75 and 0.84, respectively under DESeq), whereas HC-E, MNM, SOM, and K-means worsened. In particular, under Med scaling, K-means plummeted from 0.37 to 0.08, and HC-E from 0.40 to 0.11.

The effects of RUVs and RUVg were consistent with earlier scenarios, continuing to suppress clustering. In contrast, RUVr caused less deterioration and occasionally improved ARIs, a pattern examined next.

#### Paradoxical performance of RUVr with stronger signals

We observed that RUVr performed better as the underlying biological signal became weaker. In RUVr, the “r” denotes “residuals”: the method first fits a regression to preserve variation associated with a specified group variable, then applies factor analysis to the residuals to estimate and remove unwanted variation. In clustering, however, the group variable is unknown at the time of harmonization. Consequently, RUVr performs the regression without adjusting for the underlying sample clusters and can inadvertently treat cluster-defining signals as unwanted variation. The stronger the biological signal, the more effectively RUVr removes it, impairing cluster discovery. To test this mechanism, we reanalyzed the data using a design matrix that included the true group labels; under this supervised setting, RUVr outperformed many other methods (Fig. S2). These findings indicate that RUVr should be used with caution in clustering, particularly when good cluster separation is anticipated.

**In summary**, scaling methods and QN can modestly improve clustering for otherwise under-performing techniques when cluster separation is good, even in the absence of artifacts. However, when separation is poor, scaling and QN generally degraded performance across mid- and low-tier techniques, notably K-means and HC-E. RUVs slightly reduced performance for most techniques, while RUVg pushed all techniques toward chance. By contrast, RUVr had mixed effects on the three HC variants, and negative to negligible effect on the other techniques as cluster separation narrowed.

### Harmonization can often partially mitigate artifacts-induced losses in clustering accuracy

To assess how artifacts affect clustering and the extent to which harmonization can offset them, we analyzed augmented test datasets at three artifact magnitudes (*d* = 1.2, 1.5, and 2.0) across four signal strengths (*c* = 0.6, 1.0, 1.5, and 2.0). Introducing artifacts led to striking declines in clustering performance (Fig. 4; Table S2). Although relative trends among harmonization methods and clustering techniques generally paralleled those in artifact-free data at comparable signal strengths, absolute ARIs were often substantially lower. Harmonization sometimes attenuated the adverse effects of artifacts, but only occasionally restored clustering performance to pre-artifact heights (typically under weak artifacts and with more effective methods such as DESeq and TMM).

**Fig. 4.**
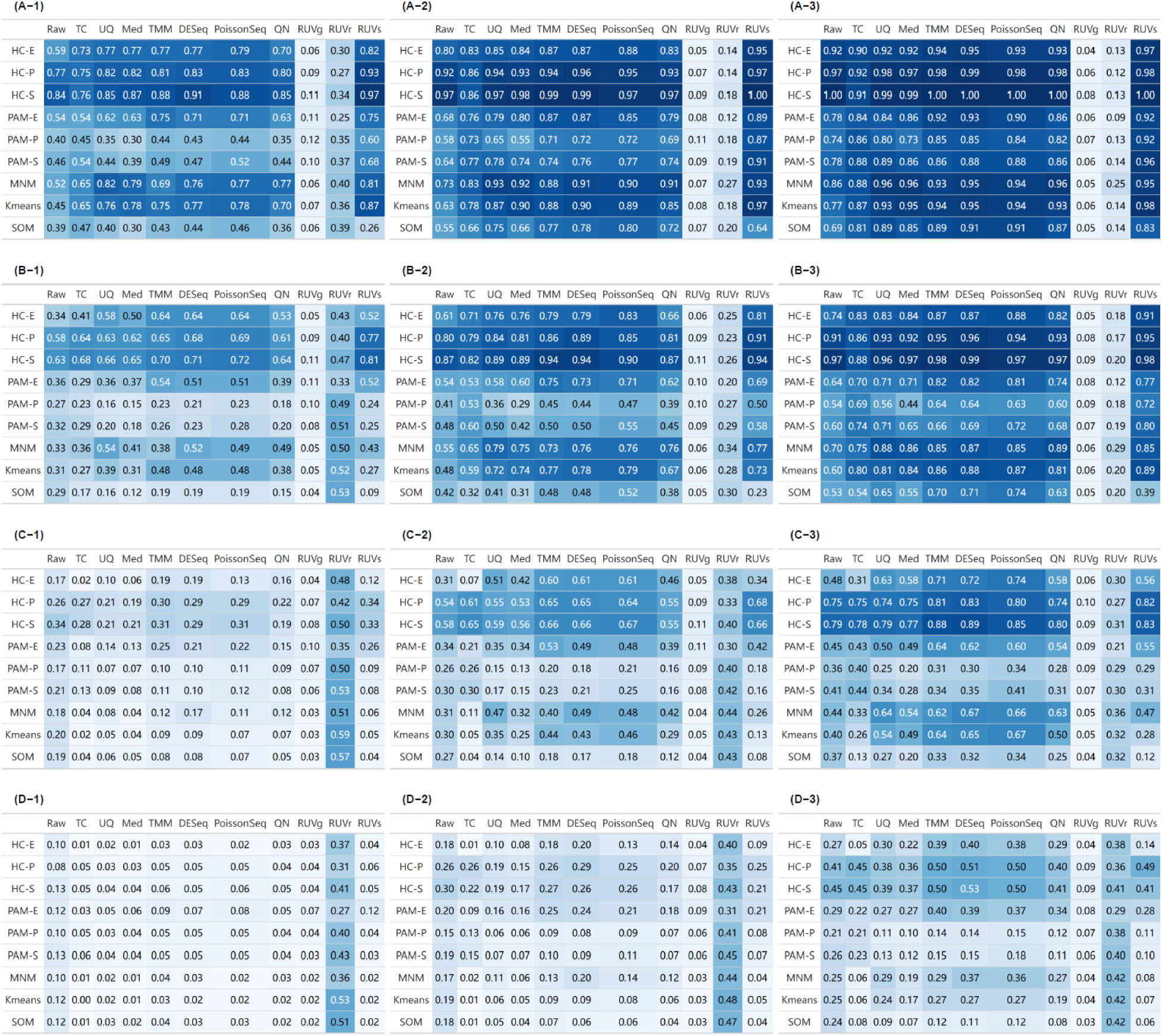
Evaluation of harmonization performance for clustering using augmented data with artifacts. Heatmaps display mean ARI values for different harmonization methods combined with various clustering techniques across four levels of biological signal strength – (A) c = 2.0; (B) c = 1.5; (C) c = 1.0; (D) c = 0.6 – and three levels of technical artifact magnitude – (1) d = 2.0; (2) d = 1.5; (3) d = 1.2. Higher ARI values (darker blue) indicate better clustering accuracy.

In the most favorable artifact-containing scenario (very strong signal, weak artifacts: *c* = 2.0, *d* = 1.2; Fig. 4A-3), both absolute ARIs and relative rankings of methods and techniques closely mirrored the moderate-signal, artifact-free scenario (Fig. 3A). The top combinations – DESeq or TMM paired with HC-P or HC- S – achieved ARIs of 0.98–1.00 (vs. 0.97–1.00 before harmonization). Runner-ups, HC-E and MNM, began around 0.92 and 0.86 and were lifted to 0.93–0.95 by DESeq or TMM. PAM-E, K-means, and SOM showed the largest recoveries (∼0.14–0.22): from 0.78, 0.77, and 0.69 pre-harmonization to 0.93 (0.92), 0.95 (0.94), and 0.91 (0.89) after DESeq (TMM). PAM-P and PAM-S also improved by ∼0.10 from 0.74 and 0.78.

#### Impact of artifact magnitude and signal strength

Performance improved as signal strengthened or artifact weakened. Overall trends across signal levels paralleled those from the benchmark data. Reducing signal to *c* = 1.5 at *d* = 1.2 decreased ARIs by ∼0.03–0.20 across techniques. Harmonization tempered these losses for better-performing methods (HC, K-means, MNM), narrowing the gaps to 0.01–0.08, but slightly widened them for worse-performers (PAM, SOM) to 0.11–0.21 (Fig. 4B-3). With even lower signal (*c* = 1.0, *d* = 1.2), cluster structure was further obscured: even the best combinations (HC paired with DESeq or TMM) yielded ARIs between 0.80 and 0.90 (Fig. 4C-3); runners-up (K-means, MNM, and PAM-E) remained ∼0.60–0.75 after harmonization, whereas others (PAM-P, PAM-S, and SOM) typically dropped below 0.45. Increasing artifact magnitude compounded losses across the board: even the top combinations (except RUVr) dropped to ∼0.65 at *d* = 1.5 (Fig. 4C-2) and to ∼0.30 at *d* = 2.0 (Fig. 4C-1), when *c* = 1.0.

#### Performance assessment of harmonization and clustering approaches

Two scaling-based methods, DESeq and TMM, consistently emerged as the top performers. Both estimate sample-specific scaling factors using robust statistics: TMM via a trimmed mean of log-fold changes and DESeq via the median of per-gene ratios. PoissonSeq also performed well, particularly with HC, though its use may be limited by the lack of ongoing maintenance of its R package.

Among regression-based methods, RUVr outperformed its peers when the signal-to-artifact ratio was low. For example, at *c* = 0.6 and *d* = 2.0, RUVr reached ARIs around 0.35, while other methods often fell below 0.10 (Fig. 4D-1). This counterintuitive result aligns with RUVr’s behavior in the artifact-free analyses. In contrast, RUVg consistently performed poorly, often yielding ARIs below 0.10. These findings held in additional sensitivity analyses for the RUV methods, varying the number of latent factors and the proportion of abundant markers used for factor estimation (Fig. S3). QN occasionally boosted performance for methods such as HC and MNM, but it rarely surpassed DESeq or TMM.

Across simulation scenarios, HC-P and HC-S, especially when paired with DESeq or TMM, remained the most effective clustering techniques, echoing the artifact-free setting. HC-E tended to sit in a second tier, with diminished ARIs under moderate to strong artifacts. PAM-E sometimes outperformed PAM-S and PAM-P, but rarely matched the ARIs of correlation-based HC. K-means, PAM, and SOM were occasionally competitive when signals were strong (*c* ≥ 1.5) and artifacts weak (*d* = 1.2), but their overall performances were more variable and generally less competitive.

**To summarize**, harmonization often partially – and occasionally fully – counteracted artifact effects, with DESeq and TMM mitigating them more effectively than alternatives.

### Harmonization is minimally sensitive to sample size, but highly sensitive to cluster-size imbalance and cluster-number misspecification

Beyond signal strength and artifact magnitude, we examined the impact of sample size, cluster-size imbalance, and cluster-number misspecification on harmonization performance in sample clustering, focusing on the strong-signal and weak-artifact scenario (*c* = 1.5, *d* = 1.2). The results showed that, while larger sample sizes offered incremental gains, both cluster-size imbalance and over-partitioning markedly degraded clustering accuracy. Notably, MNM performed comparatively robust under these conditions.

#### Sample size variation

We evaluated datasets with progressively smaller per-cluster sizes (50 and 27 samples; Fig. S4). ARIs declined modestly as sample size decreased, but relative methods rankings were stable. This finding supports prior reports that clustering performance depends more on data quality and intrinsic structure than on sample size^44^.

#### Cluster-size imbalance

Imbalance, defined as the MXF:UPS sample-size ratio, exerted a pronounced impact (Fig. S5). Under severe imbalance (1:9), most combinations of harmonization methods and clustering techniques yielded ARIs near zero (<0.10). A notable exception was RUVr paired with MNM, reaching ARIs around 0.77. As balance improved, ARIs rose across methods and RUVr’s advantage diminished. DESeq and TMM typically outpaced RUVr near parity, indicating that while RUVr plus MNM can mitigate extreme imbalance, DESeq or TMM are preferable when groups are similar in size.

#### Cluster-number misspecification

As expected, over-partitioning (i.e., specifying more clusters than truly present) broadly degraded ARIs (Fig. S6). HC-P and HC-S were particularly sensitive: when combined with DESeq or TMM, ARIs fell sharply from >0.95 at K=2 to ∼0.55 at K=4, declined further to ∼0.37 at K=6, and dropped below 0.30 at K=8. HC-E exhibited similar sensitivity, with ARIs lowering from 0.88 to 0.47 when paired with TMM. By contrast, MNM was more resilient, with ARIs decreasing only modestly from 0.85 to 0.60. These results suggest that correlation-based distances amplify the adverse effects of a misspecified K, whereas model-based approaches like MNM offer greater robustness.

**In aggregate**, Fig. 5 summarizes the overall performance patterns across these data-generation settings.

**Fig. 5.**
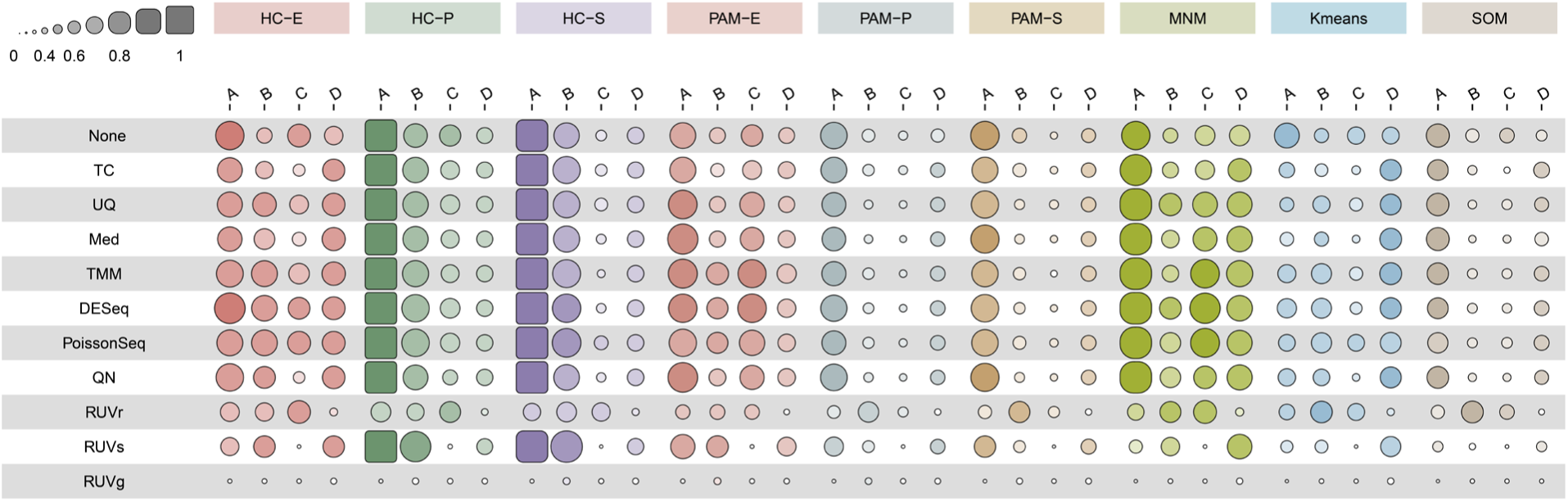
Summary of harmonization performance in clustering across data-generation settings for the augmented data. The dot-matrix plot depicts the mean ARI for combinations of harmonization methods (rows) and clustering techniques (column groups) across four representative scenarios, each drawn from a distinct data-generation setting: (A) weak signal (c = 0.6) without artifacts; (B) strong signal (c = 1.5) with strong artifacts (d = 2.0); (C) cluster-size imbalance (MXF:PMFH = 1:4); and (D) cluster-number misspecification (K = 6). Scenarios A and B use equal cluster sizes (1:1) and specified the correct number of clusters (K = 2); scenarios C and D use strong signal (*c* = 1.5) and weak artifacts (*d* = 1.2); in addition, scenario C specifies the correct number of true clusters (K = 2), while scenario D uses equal cluster sizes (1:1); all four scenarios keep the total sample size at 200. Circle size and color intensity are proportional to ARI values.

### TCGA breast cancer case study confirms the top-performing harmonization methods

In the TCGA breast cancer cohort, miRNA sequencing depth rose steadily from 2010 through August 2011 and then plateaued, providing clear evidence of time-related handling artifacts (Fig. S7). The Spearman correlation between sequencing depth and data collection date was 0.68 (p < 0.001). Strong biological differences were also observed: 29% of markers (550/1,881) were differentially expressed (adjusted p < 0.01) between Luminal A and Basal tumors using TC-scaled data.

Across the four artifact scenarios for cohort reconstruction, results broadly mirrored the simulations. DESeq and TMM remained the most effective overall (Fig. 6). TC performed better in TCGA than in augmented MSK datasets, ranking among the top methods in three of the four scenarios across most clustering techniques. By contrast, UQ and Med were substantially weaker, especially in Scenario I (weak artifacts) and Scenario II (strong, balanced artifacts). Also consistent with the simulations, HC-P and HC-S stayed the best techniques, while PAM-P and PAM-S were the weakest.

**Fig. 6.**
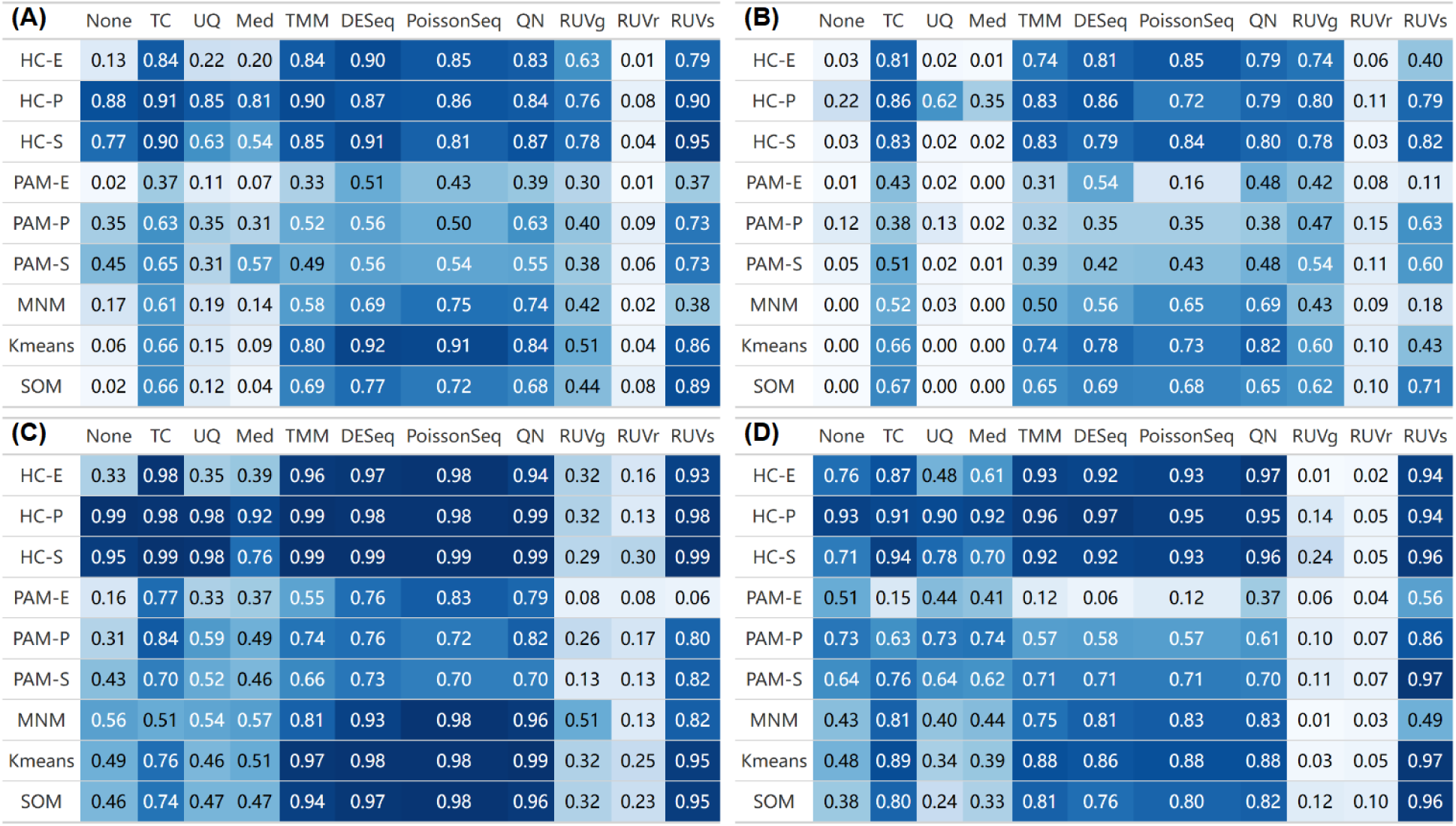
Harmonization–clustering performance on miRNA sequencing from reconstructed TCGA breast cancer cohorts. (A) Scenario I: weak artifacts (all samples from the late batch). (B) Scenario II: strong, balanced artifacts (samples drawn equally from the early and late batches). (C) Scenario III: strong, confounded artifacts (Luminal A samples from the late batch and Basal samples from the early batch). (D) Scenario IV: strong, confounded artifacts in the opposite direction (Luminal A samples from the early batch and Basal samples from the late batch).

Interestingly, Scenario II produced poorer clustering than Scenario I, whereas Scenarios III and IV (strong, confounded artifacts) showed improved performance. This pattern suggests that artifacts highly correlated (either positively or negatively) with the true subtypes can artificially enhance cluster separability, albeit with biased cluster centers. Such shifts in artifact-signal confounding provide a mechanistic explanation for the lack of replicability often observed in practice^45,46^.

## Discussion

In this study, we systematically evaluated multiple harmonization methods and their interactions with clustering techniques for miRNA sequencing data. Using deep generative models, we created datasets that captured the distributional complexity of empirical data, enabling systematic benchmarking across signal-to-artifact ratios and cluster-composition settings. To complement the simulations, we constructed cohorts with varying artifact patterns from publicly available TCGA data, providing additional real-world validation.

Our results highlight the critical role of harmonization in improving clustering accuracy, especially at moderate signal-to-artifact ratios. As expected, harmonization yielded limited gains when the ratios were very high (little to fix) or very low (insufficient rescue). Beyond signal strength and artifact magnitude, performance was strongly affected by cluster-size imbalance and cluster-number misspecification, both of which can substantially impair clustering, though robust harmonization and model-based clustering can partly mitigate these effects.

No single method was universally optimal. However, two scaling methods, DESeq and TMM, consistently ranked among the top performers across nearly all simulation settings and reconstructed cohort scenarios. RUVr was particularly useful when the signal-to-artifact ratio was low or the cluster composition was highly imbalanced, whereas RUVg consistently degraded the clustering, likely reflecting the difficulty of identifying suitable negative-control markers in miRNA data^32^. Among clustering techniques, correlation-based hierarchical clustering generally outperformed alternatives when the number of clusters was specified correctly, while model-based approaches like MNM were more robust to cluster-size imbalance and cluster-number misspecification. Because the empirical data lacked an explicit batch variable, we did not assess batch-effect correction methods such as ComBat^26^.

Overall, our findings underscore the importance of jointly tailoring harmonization and clustering strategies to the characteristics of each dataset. By systematic comparison across diverse scenarios, we provide a practical framework for evaluating these strategies in miRNA clustering studies, thereby supporting the development of more effective analytical pipelines and informing best practices. Because sample clustering can reveal novel subgroups and refine disease subtyping, adopting such evidence-based practices will yield more reliable biological insights and ultimately advance disease management.

## Supporting information

Full Supplement

## Data Availability

The empirical and simulated miRNA sequencing dataset pairs used in this study are available from the *PRECISION.seq.augmented* data release: https://github.com/Omics-Data-Harmonization-EBP/PRECISION.seq.augmented/releases/download/Data/MSKpair_augmented.rds. The TCGA breast cancer miRNA sequencing data are publicly accessible via the Genomic Data Commons: https://portal.gdc.cancer.gov/. Batch annotations for TCGA breast cancer data were derived using the *MBatch* package, and subtype labels were obtained from the supplementary materials of Ellrott et al. (2025).

## Code Availability

The *PRECISION.seq.augmented* R package is openly available on GitHub: https://github.com/Omics-Data-Harmonization-EBP/PRECISION.seq.augmented. Reproducible R scripts for this manuscript, including the analysis of the TCGA breast cancer data and generation of all figures, are provided also on GitHub: https://github.com/Omics-Data-Harmonization-EBP/PRECISION.seq.augmented/tree/main/paper-supplementary-materials.

## Acknowledgements

We thank Professor Thomas Tuschl for assistance with sequencing the sarcoma samples and Terry Helms for graphic design support on Figure 1.

## Author Contributions

**Conceptualization**: LXQ, JZ, YD, YW, YX, XW, and YQ; **Methodology**: LXQ, JZ, YD, YW, YX, XW, and YQ; **Data Curation**: JZ, YD, YW, YX, XW, YQ, MW, SS, and LXQ; **Formal Analysis**: JZ, YD, YW, YX, XW, YQ, and LXQ; **Writing – Original Draft**: JZ and LXQ; **Writing – Review and Editing**: all authors; **Resources**: LXQ; **Supervision**: LXQ; **Funding Acquisition**: LXQ and SS

## Key Points

- The effectiveness of sequencing data harmonization depends on the downstream analytic task and should therefore be evaluated in an analysis-specific context.
- We systematically evaluated harmonization methods for microRNA sequencing in the context of sample clustering using controlled, realistic data-generation regimes that spandiverse scenarios of signal-to-artifact ratio, sample size, cluster-size imbalance, and cluster-number misspecification.
- An open-source R package implementing this contextual evaluation framework for microRNA sequencing data harmonization is available on GitHub.

## Table of Abbreviations

ARI: Adjusted Rand Index
BRCA: Breast Cancer (TCGA study abbreviation)
cARI: Complementary Adjusted Rand Index
HC-E: Hierarchical Clustering with Euclidean distance
HC-P: Hierarchical Clustering with Pearson correlation
HC-S: Hierarchical Clustering with Spearman correlation
IQR: Inter-Quartile Range
Med: Median Normalization
miRNA: MicroRNA
MNM: Multivariate Normal Mixture
MXF: Myxofibrosarcoma
PAM-E: Partitioning Around Medoids with Euclidean distance
PAM-P: Partitioning Around Medoids with Pearson correlation
PAM-S: Partitioning Around Medoids with Spearman correlation
PoissonSeq: Poisson Based Normalization
QN: Quantile Normalization
RUVg: Remove Unwanted Variation using negative control markers
RUVr: Remove Unwanted Variation using residuals
RUVs: Remove Unwanted Variation using replicate samples
SOM: Self-Organizing Map
TC: Total Count Normalization
TCGA: The Cancer Genome Atlas
TMM: Trimmed Mean of M-values Normalization
UMAP: Uniform Manifold Approximation and Projection
UPS: Undifferentiated Pleomorphic Sarcoma
UQ: Upper Quartile Normalization

## References

1. Bartel, D. P. MicroRNAs: genomics, biogenesis, mechanism, and function. Cell 116, 281–297 (2004).

2. Ambros, V. The functions of animal microRNAs. Nature 431, 350–355 (2004).

3. Cummins, J. M. & Velculescu, V. E. Implications of micro-RNA profiling for cancer diagnosis. Oncogene 25, 6220–6227 (2006).

4. Cantini, L. et al. Identification of microRNA clusters cooperatively acting on epithelial to mesenchymal transition in triple negative breast cancer. Nucleic Acids Res. 47, 2205–2215 (2019).

5. Bullard, J. H., Purdom, E., Hansen, K. D. & Dudoit, S. Evaluation of statistical methods for normalization and differential expression in mRNA-Seq experiments. BMC Bioinformatics 11, 94 (2010).

6. Sahoo, O. S. et al. Role of next□generation sequencing in revolutionizing healthcare for cancer management. MedComm Futur. Med. 3, (2024).

7. Dillies, M.-A. et al. A comprehensive evaluation of normalization methods for Illumina high-throughput RNA sequencing data analysis. Brief. Bioinform. 14, 671–683 (2013).

8. Qin, L.-X. et al. Statistical assessment of depth normalization for small RNA sequencing. *JCO Clin*. Cancer Inform. 4, 567–582 (2020).

9. Zou, J., Düren, Y. & Qin, L.-X. PRECISION.Seq: An R package for benchmarking depth normalization in microRNA sequencing. Front. Genet. 12, 823431 (2021).

10. Babak, T., Zhang, W., Morris, Q., Blencowe, B. J. & Hughes, T. R. Probing microRNAs with microarrays: tissue specificity and functional inference. RNA 10, 1813–1819 (2004).

11. Subramanian, S. et al. MicroRNA expression signature of human sarcomas. Oncogene 27, 2015– 2026 (2008).

12. Ludwig, N. et al. Distribution of miRNA expression across human tissues. Nucleic Acids Res. 44, 3865–3877 (2016).

13. Chu, A. et al. Large-scale profiling of microRNAs for The Cancer Genome Atlas. Nucleic Acids Res. 44, e3 (2016).

14. Qin, L.-X., Huang, H.-C. & Begg, C. B. Cautionary note on using cross-validation for molecular classification. J. Clin. Oncol. 34, 3931–3938 (2016).

15. Huang, H.-C. & Qin, L.-X. Empirical evaluation of data normalization methods for molecular classification. PeerJ 6, e4584 (2018).

16. Wu, Y., Huang, H.-C. & Qin, L.-X. Making external validation valid for molecular classifier development. *JCO Precis*. Oncol. 5, 1250–1258 (2021).

17. Huang, H.-C., Wu, Y., Yang, Q. & Qin, L.-X. PRECISION.Array: An R package for benchmarking microRNA array data normalization in the context of sample classification. Front. Genet. 13, 838679 (2022).

18. Wu, Y., Yuen, B. W.-Y., Wei, Y. & Qin, L.-X. On data normalization and batch-effect correction for tumor subtyping with microRNA data. NAR Genom. Bioinform. 5, lqac100 (2023).

19. Ni, A. & Qin, L.-X. Performance evaluation of transcriptomics data normalization for survival risk prediction. *arXiv [q-bio.GN]* bbab257 (2021) doi:10.1093/bib/bbab257.

20. Qi, Y., Wang, X. & Qin, L.-X. Optimizing sample size for supervised machine learning with bulk transcriptomic sequencing: a learning curve approach. Brief. Bioinform. 26, bbaf097 (2025).

21. Hutter, C. & Zenklusen, J. C. The Cancer Genome Atlas: Creating lasting value beyond its data. Cell 173, 283–285 (2018).

22. Lee, A. Y. et al. Optimal percent myxoid component to predict outcome in high-grade myxofibrosarcoma and undifferentiated pleomorphic sarcoma. Ann. Surg. Oncol. 23, 818–825 (2016).

23. McInnes, L., Healy, J. & Melville, J. UMAP: Uniform Manifold Approximation and Projection for Dimension Reduction. *arXiv [stat.ML]* (2018).

24. Kerr, M. K., Martin, M. & Churchill, G. A. Analysis of variance for gene expression microarray data. J. Comput. Biol. 7, 819–837 (2000).

25. Qin, L.-X. & Satagopan, J. M. Normalization method for transcriptional studies of heterogeneous samples--simultaneous array normalization and identification of equivalent expression. Stat. Appl. Genet. Mol. Biol. 8, Article 10 (2009).

26. Zhang, Y., Parmigiani, G. & Johnson, W. E. ComBat-seq: batch effect adjustment for RNA-seq count data. NAR Genom. Bioinform. 2, lqaa078 (2020).

27. Cancer Genome Atlas Research Network et al. The Cancer Genome Atlas Pan-Cancer analysis project. Nat. Genet. 45, 1113–1120 (2013).

28. Robinson, M. D. & Smyth, G. K. Moderated statistical tests for assessing differences in tag abundance. Bioinformatics 23, 2881–2887 (2007).

29. Anders, S. & Huber, W. Differential expression analysis for sequence count data. Genome Biol. 11, R106 (2010).

30. Li, J., Witten, D. M., Johnstone, I. M. & Tibshirani, R. Normalization, testing, and false discovery rate estimation for RNA-sequencing data. Biostatistics 13, 523–538 (2012).

31. Bolstad, B. M., Irizarry, R. A., Astrand, M. & Speed, T. P. A comparison of normalization methods for high density oligonucleotide array data based on variance and bias. Bioinformatics 19, 185–193 (2003).

32. Risso, D., Ngai, J., Speed, T. P. & Dudoit, S. Normalization of RNA-seq data using factor analysis of control genes or samples. Nat. Biotechnol. 32, 896–902 (2014).

33. Forgy, E. W. Cluster analysis of multivariate data : efficiency versus interpretability of classifications. Biometrics 21, 768–769 (1965).

34. Kaufman, L. & Rousseeuw, P. J. Finding groups in data: an introduction to cluster analysis. (2009).

35. Kohonen, T. The self-organizing map. Proc. IEEE (2002).

36. Fraley, C. & Raftery, A. E. Model-based clustering, discriminant analysis, and density estimation. J. Am. Stat. Assoc. 97, 611–631 (2002).

37. Anderberg, M. R. Cluster analysis for applications: probability and mathematical statistics: a series of monographs and textbooks. 19, (2014).

38. Scrucca, L., Fraley, C., Murphy, T. B. & Adrian E., R. Model-Based Clustering, Classification, and Density Estimation Using Mclust in R. (Chapman and Hall/CRC, Boca Raton, 2023).

39. Rand, W. M. Objective criteria for the evaluation of clustering methods. J. Am. Stat. Assoc. 66, 846– 850 (1971).

40. Hubert, L. & Arabie, P. Comparing partitions. J. Classif. 2, 193–218 (1985).

41. Gao, G. F. et al. Before and after: Comparison of legacy and harmonized TCGA Genomic Data Commons’ data. Cell Syst. 9, 24–34.e10 (2019).

42. Colaprico, A. et al. TCGAbiolinks: an R/Bioconductor package for integrative analysis of TCGA data. Nucleic Acids Res. 44, e71 (2016).

43. Ellrott, K. et al. Classification of non-TCGA cancer samples to TCGA molecular subtypes using compact feature sets. Cancer Cell 43, 195–212.e11 (2025).

44. Vidman, L., Källberg, D. & Rydén, P. Cluster analysis on high dimensional RNA-seq data with applications to cancer research - An evaluation study. PLoS One 14, e0219102 (2019).

45. Guinney, J. et al. The consensus molecular subtypes of colorectal cancer. Nature medicine 21, (2015).

46. Aran, D., Sirota, M. & Butte, A. J. Systematic pan-cancer analysis of tumour purity. Nat. Commun. 6, 8971 (2015).

