## Supplementary material for "Contextual Evaluation of MicroRNA Sequencing Data Harmonization: Performance in Sample Clustering": Full Supplement

**Supplementary Figures: S1-S6**

**Supplementary Table: S1**

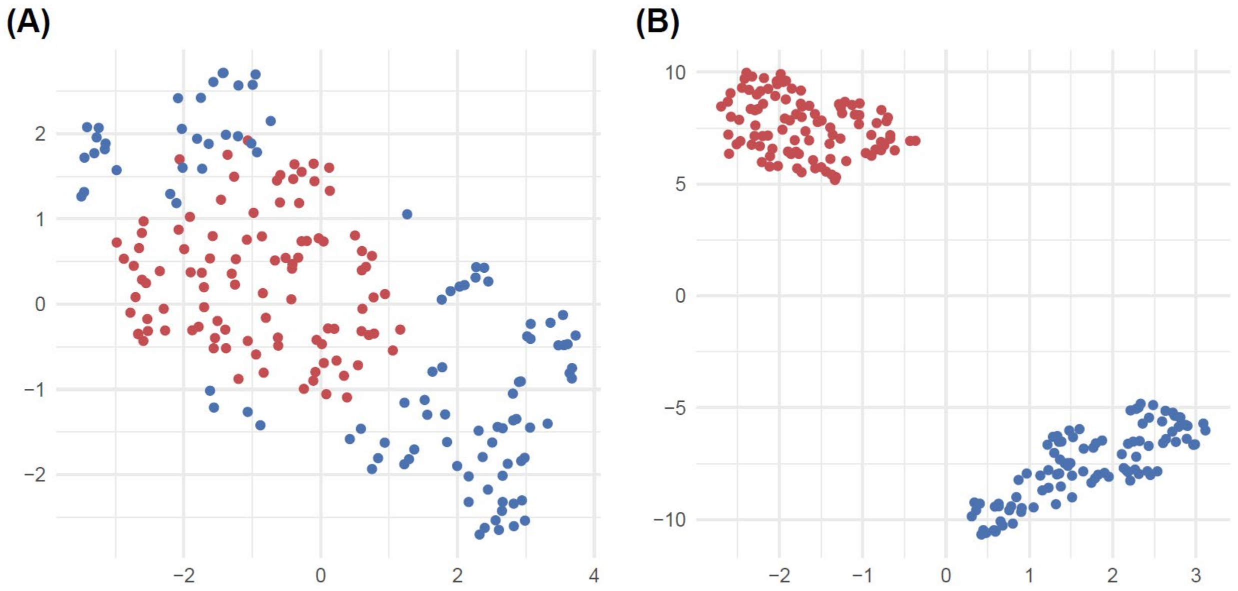


**Fig. S1 | Harmonization performance of RUVg vs. TMM in the absence of technical artifacts.** UMAPs of a randomly selected dataset generated with moderate biological signal (c=1.0) and no technical artifacts, after harmonization using (A) RUVg and (B) TMM. Colors indicate sarcoma subtypes: MXF (red) and UPS (blue).

(A)

| | Raw | TC | UQ | Med | TMM | DESeq | PoissonSeq | QN | RUVg | RUVr | RUVs |
|---|---|---|---|---|---|---|---|---|---|---|---|
| HC-E | 0.96 | 0.97 | 0.99 | 0.98 | 0.99 | 1.00 | 1.00 | 0.99 | 0.03 | 1.00 | 0.88 |
| HC-P | 1.00 | 0.99 | 1.00 | 1.00 | 1.00 | 1.00 | 1.00 | 1.00 | 0.03 | 1.00 | 1.00 |
| HC-S | 1.00 | 0.99 | 1.00 | 1.00 | 1.00 | 1.00 | 1.00 | 1.00 | 0.03 | 1.00 | 1.00 |
| PAM-E | 0.81 | 0.82 | 0.87 | 0.90 | 0.88 | 0.89 | 0.84 | 0.90 | 0.03 | 0.98 | 0.80 |
| PAM-P | 0.89 | 0.87 | 0.79 | 0.80 | 0.85 | 0.87 | 0.83 | 0.87 | 0.02 | 0.97 | 0.80 |
| PAM-S | 0.92 | 0.88 | 0.87 | 0.89 | 0.89 | 0.90 | 0.87 | 0.89 | 0.02 | 0.99 | 0.86 |
| MNM | 0.89 | 0.93 | 0.99 | 0.98 | 0.94 | 0.95 | 0.93 | 0.94 | 0.01 | 0.98 | 0.74 |
| Kmeans | 0.83 | 0.85 | 0.88 | 0.93 | 0.95 | 0.95 | 0.87 | 0.90 | 0.00 | 1.00 | 0.68 |
| SOM | 0.75 | 0.78 | 0.87 | 0.90 | 0.88 | 0.86 | 0.83 | 0.85 | 0.01 | 0.99 | 0.39 |

(B)

| | Raw | TC | UQ | Med | TMM | DESeq | PoissonSeq | QN | RUVg | RUVr | RUVs |
|---|---|---|---|---|---|---|---|---|---|---|---|
| HC-E | 0.71 | 0.61 | 0.62 | 0.60 | 0.67 | 0.78 | 0.64 | 0.69 | 0.04 | 1.00 | 0.42 |
| HC-P | 0.97 | 0.99 | 0.99 | 0.99 | 0.99 | 1.00 | 1.00 | 0.99 | 0.04 | 1.00 | 0.99 |
| HC-S | 0.94 | 0.98 | 0.98 | 0.98 | 0.99 | 0.99 | 0.99 | 0.98 | 0.04 | 1.00 | 0.93 |
| PAM-E | 0.64 | 0.65 | 0.72 | 0.76 | 0.73 | 0.71 | 0.68 | 0.75 | 0.05 | 0.93 | 0.60 |
| PAM-P | 0.66 | 0.64 | 0.59 | 0.58 | 0.59 | 0.65 | 0.59 | 0.66 | 0.03 | 0.87 | 0.45 |
| PAM-S | 0.72 | 0.64 | 0.68 | 0.71 | 0.65 | 0.69 | 0.66 | 0.71 | 0.03 | 0.93 | 0.52 |
| MNM | 0.71 | 0.80 | 0.84 | 0.83 | 0.83 | 0.84 | 0.82 | 0.84 | 0.01 | 0.94 | 0.28 |
| Kmeans | 0.61 | 0.35 | 0.31 | 0.30 | 0.42 | 0.46 | 0.38 | 0.40 | 0.01 | 0.99 | 0.25 |
| SOM | 0.57 | 0.50 | 0.55 | 0.56 | 0.52 | 0.50 | 0.49 | 0.53 | 0.01 | 0.95 | 0.21 |

(C)

| | Raw | TC | UQ | Med | TMM | DESeq | PoissonSeq | QN | RUVg | RUVr | RUVs |
|---|---|---|---|---|---|---|---|---|---|---|---|
| HC-E | 0.40 | 0.16 | 0.14 | 0.11 | 0.16 | 0.19 | 0.16 | 0.17 | 0.05 | 0.98 | 0.17 |
| HC-P | 0.69 | 0.80 | 0.70 | 0.80 | 0.74 | 0.84 | 0.68 | 0.77 | 0.06 | 0.99 | 0.65 |
| HC-S | 0.65 | 0.65 | 0.58 | 0.67 | 0.66 | 0.74 | 0.60 | 0.67 | 0.05 | 0.98 | 0.33 |
| PAM-E | 0.41 | 0.37 | 0.50 | 0.52 | 0.44 | 0.44 | 0.40 | 0.45 | 0.07 | 0.79 | 0.29 |
| PAM-P | 0.28 | 0.32 | 0.34 | 0.34 | 0.27 | 0.31 | 0.29 | 0.37 | 0.04 | 0.61 | 0.15 |
| PAM-S | 0.35 | 0.28 | 0.41 | 0.42 | 0.28 | 0.29 | 0.31 | 0.38 | 0.04 | 0.73 | 0.18 |
| MNM | 0.43 | 0.25 | 0.18 | 0.14 | 0.33 | 0.39 | 0.29 | 0.31 | 0.01 | 0.79 | 0.07 |
| Kmeans | 0.37 | 0.16 | 0.10 | 0.09 | 0.15 | 0.16 | 0.14 | 0.15 | 0.02 | 0.89 | 0.11 |
| SOM | 0.36 | 0.26 | 0.26 | 0.27 | 0.19 | 0.19 | 0.18 | 0.20 | 0.02 | 0.76 | 0.10 |

**Fig. S2 | Harmonization performance in the absence of technical artifacts, when sample group labels are provided to RUVr.** Heatmaps show the mean Adjusted Rand Index (ARI) at three signal levels: (A) c = 1.0, (B) c = 0.6, and (C) c = 0.2. Darker blue denotes higher ARI (better clustering accuracy). In this supervised setting, RUVr outperforms many other harmonization methods.

| k = 1, proportion = 0.50 | RUVg | RUVr | RUVs |
|---|---|---|---|
| HC-E | 0.05 | 0.18 | 0.91 |
| HC-P | 0.07 | 0.17 | 0.95 |
| HC-S | 0.08 | 0.19 | 0.98 |
| PAM-E | 0.07 | 0.12 | 0.80 |
| PAM-P | 0.08 | 0.18 | 0.71 |
| PAM-S | 0.07 | 0.19 | 0.80 |
| MNM | 0.05 | 0.27 | 0.86 |
| Kmeans | 0.05 | 0.20 | 0.90 |
| SOM | 0.04 | 0.20 | 0.38 |

| k = 1, proportion = 0.75 | RUVg | RUVr | RUVs |
|---|---|---|---|
| HC-E | 0.05 | 0.19 | 0.92 |
| HC-P | 0.07 | 0.17 | 0.96 |
| HC-S | 0.09 | 0.20 | 0.99 |
| PAM-E | 0.07 | 0.13 | 0.79 |
| PAM-P | 0.09 | 0.19 | 0.72 |
| PAM-S | 0.07 | 0.20 | 0.80 |
| MNM | 0.06 | 0.29 | 0.85 |
| Kmeans | 0.06 | 0.21 | 0.88 |
| SOM | 0.05 | 0.21 | 0.39 |

| k = 1, proportion = 0.90 | RUVg | RUVr | RUVs |
|---|---|---|---|
| HC-E | 0.05 | 0.19 | 0.92 |
| HC-P | 0.08 | 0.18 | 0.95 |
| HC-S | 0.09 | 0.21 | 0.98 |
| PAM-E | 0.07 | 0.13 | 0.79 |
| PAM-P | 0.09 | 0.19 | 0.72 |
| PAM-S | 0.08 | 0.20 | 0.79 |
| MNM | 0.06 | 0.30 | 0.86 |
| Kmeans | 0.06 | 0.21 | 0.89 |
| SOM | 0.05 | 0.21 | 0.38 |

| k = 3, proportion = 0.50 | RUVg | RUVr | RUVs |
|---|---|---|---|
| HC-E | 0.00 | 0.01 | 0.97 |
| HC-P | 0.01 | 0.01 | 0.97 |
| HC-S | 0.01 | 0.01 | 1.00 |
| PAM-E | 0.01 | 0.01 | 0.94 |
| PAM-P | 0.02 | 0.03 | 0.94 |
| PAM-S | 0.02 | 0.03 | 0.98 |
| MNM | 0.00 | 0.04 | 0.96 |
| Kmeans | 0.02 | 0.03 | 1.00 |
| SOM | 0.02 | 0.03 | 0.99 |

| k = 3, proportion = 0.75 | RUVg | RUVr | RUVs |
|---|---|---|---|
| HC-E | 0.01 | 0.01 | 0.97 |
| HC-P | 0.01 | 0.01 | 0.97 |
| HC-S | 0.01 | 0.02 | 1.00 |
| PAM-E | 0.01 | 0.01 | 0.94 |
| PAM-P | 0.03 | 0.03 | 0.94 |
| PAM-S | 0.03 | 0.03 | 0.98 |
| MNM | 0.00 | 0.05 | 0.96 |
| Kmeans | 0.02 | 0.03 | 1.00 |
| SOM | 0.03 | 0.04 | 0.99 |

| k = 3, proportion = 0.90 | RUVg | RUVr | RUVs |
|---|---|---|---|
| HC-E | 0.01 | 0.01 | 0.97 |
| HC-P | 0.01 | 0.01 | 0.97 |
| HC-S | 0.01 | 0.01 | 1.00 |
| PAM-E | 0.01 | 0.01 | 0.94 |
| PAM-P | 0.03 | 0.03 | 0.94 |
| PAM-S | 0.02 | 0.03 | 0.98 |
| MNM | 0.00 | 0.04 | 0.96 |
| Kmeans | 0.02 | 0.03 | 1.00 |
| SOM | 0.03 | 0.03 | 0.99 |

| k = 5, proportion = 0.50 | RUVg | RUVr | RUVs |
|---|---|---|---|
| HC-E | 0.00 | 0.00 | 0.98 |
| HC-P | 0.01 | 0.01 | 0.97 |
| HC-S | 0.01 | 0.01 | 1.00 |
| PAM-E | 0.00 | 0.00 | 0.96 |
| PAM-P | 0.02 | 0.01 | 0.98 |
| PAM-S | 0.02 | 0.01 | 0.99 |
| MNM | 0.00 | 0.03 | 0.95 |
| Kmeans | 0.01 | 0.01 | 1.00 |
| SOM | 0.02 | 0.01 | 1.00 |

| k = 5, proportion = 0.75 | RUVg | RUVr | RUVs |
|---|---|---|---|
| HC-E | 0.00 | 0.00 | 0.97 |
| HC-P | 0.00 | 0.01 | 0.97 |
| HC-S | 0.00 | 0.01 | 1.00 |
| PAM-E | 0.00 | 0.00 | 0.97 |
| PAM-P | 0.01 | 0.01 | 0.98 |
| PAM-S | 0.01 | 0.01 | 0.99 |
| MNM | 0.00 | 0.03 | 0.95 |
| Kmeans | 0.01 | 0.01 | 1.00 |
| SOM | 0.02 | 0.01 | 1.00 |

| k = 5, proportion = 0.90 | RUVg | RUVr | RUVs |
|---|---|---|---|
| HC-E | 0.00 | 0.01 | 0.97 |
| HC-P | 0.00 | 0.01 | 0.97 |
| HC-S | 0.00 | 0.01 | 1.00 |
| PAM-E | 0.00 | 0.00 | 0.96 |
| PAM-P | 0.02 | 0.01 | 0.98 |
| PAM-S | 0.01 | 0.01 | 0.99 |
| MNM | 0.00 | 0.03 | 0.95 |
| Kmeans | 0.01 | 0.01 | 1.00 |
| SOM | 0.01 | 0.01 | 1.00 |

**Fig. S3 | Harmonization performance of RUV methods under alternative factor-analysis settings.** Heatmaps show the mean ARI for RUV factor modeling with varying numbers of latent factor – (top row) 1, (middle row)3, and (bottom row) 5 – and varying proportions of abundant markers used for latent factor estimation – (left column) 50%, (middle column) 75%, and (right column) 90%, under strong signal ($c$ = 1.5) and weak artifacts ($d$ = 1.2).

(A)

| | Raw | TC | UQ | Med | TMM | DESeq | PoissonSeq | QN | RUVg | RUVr | RUVs |
|---|---|---|---|---|---|---|---|---|---|---|---|
| HC-E | 0.76 | 0.85 | 0.85 | 0.82 | 0.87 | 0.88 | 0.88 | 0.84 | 0.05 | 0.18 | 0.90 |
| HC-P | 0.93 | 0.87 | 0.94 | 0.92 | 0.96 | 0.96 | 0.94 | 0.93 | 0.07 | 0.17 | 0.95 |
| HC-S | 0.97 | 0.88 | 0.96 | 0.96 | 0.99 | 0.98 | 0.96 | 0.97 | 0.09 | 0.19 | 0.99 |
| PAM-E | 0.65 | 0.70 | 0.73 | 0.74 | 0.85 | 0.83 | 0.81 | 0.76 | 0.07 | 0.13 | 0.79 |
| PAM-P | 0.55 | 0.70 | 0.55 | 0.44 | 0.64 | 0.65 | 0.65 | 0.59 | 0.08 | 0.18 | 0.70 |
| PAM-S | 0.60 | 0.74 | 0.69 | 0.64 | 0.67 | 0.69 | 0.71 | 0.67 | 0.07 | 0.19 | 0.79 |
| MNM | 0.70 | 0.76 | 0.88 | 0.86 | 0.86 | 0.87 | 0.85 | 0.89 | 0.05 | 0.28 | 0.85 |
| Kmeans | 0.60 | 0.81 | 0.81 | 0.85 | 0.87 | 0.88 | 0.88 | 0.82 | 0.06 | 0.20 | 0.88 |
| SOM | 0.54 | 0.52 | 0.66 | 0.55 | 0.71 | 0.72 | 0.74 | 0.64 | 0.04 | 0.20 | 0.38 |

(B)

| | Raw | TC | UQ | Med | TMM | DESeq | PoissonSeq | QN | RUVg | RUVr | RUVs |
|---|---|---|---|---|---|---|---|---|---|---|---|
| HC-E | 0.73 | 0.78 | 0.82 | 0.83 | 0.86 | 0.88 | 0.87 | 0.80 | 0.05 | 0.17 | 0.89 |
| HC-P | 0.91 | 0.87 | 0.91 | 0.90 | 0.92 | 0.93 | 0.91 | 0.91 | 0.07 | 0.15 | 0.95 |
| HC-S | 0.92 | 0.88 | 0.93 | 0.93 | 0.97 | 0.96 | 0.94 | 0.94 | 0.08 | 0.18 | 0.98 |
| PAM-E | 0.64 | 0.68 | 0.70 | 0.71 | 0.81 | 0.80 | 0.79 | 0.72 | 0.07 | 0.11 | 0.78 |
| PAM-P | 0.56 | 0.69 | 0.57 | 0.46 | 0.65 | 0.65 | 0.64 | 0.60 | 0.09 | 0.17 | 0.77 |
| PAM-S | 0.63 | 0.75 | 0.70 | 0.65 | 0.67 | 0.69 | 0.72 | 0.67 | 0.08 | 0.18 | 0.81 |
| MNM | 0.68 | 0.75 | 0.87 | 0.87 | 0.85 | 0.87 | 0.86 | 0.87 | 0.05 | 0.23 | 0.91 |
| Kmeans | 0.60 | 0.74 | 0.81 | 0.82 | 0.86 | 0.87 | 0.86 | 0.79 | 0.06 | 0.18 | 0.88 |
| SOM | 0.53 | 0.50 | 0.63 | 0.52 | 0.67 | 0.67 | 0.70 | 0.59 | 0.05 | 0.19 | 0.44 |

(C)

| | Raw | TC | UQ | Med | TMM | DESeq | PoissonSeq | QN | RUVg | RUVr | RUVs |
|---|---|---|---|---|---|---|---|---|---|---|---|
| HC-E | 0.67 | 0.68 | 0.83 | 0.80 | 0.85 | 0.87 | 0.87 | 0.81 | 0.05 | 0.15 | 0.86 |
| HC-P | 0.89 | 0.86 | 0.90 | 0.90 | 0.92 | 0.93 | 0.91 | 0.90 | 0.07 | 0.14 | 0.96 |
| HC-S | 0.92 | 0.88 | 0.91 | 0.90 | 0.94 | 0.94 | 0.92 | 0.91 | 0.08 | 0.16 | 0.97 |
| PAM-E | 0.63 | 0.62 | 0.68 | 0.70 | 0.78 | 0.78 | 0.75 | 0.70 | 0.06 | 0.10 | 0.78 |
| PAM-P | 0.59 | 0.68 | 0.60 | 0.53 | 0.66 | 0.66 | 0.66 | 0.62 | 0.10 | 0.16 | 0.78 |
| PAM-S | 0.64 | 0.75 | 0.72 | 0.66 | 0.69 | 0.72 | 0.73 | 0.69 | 0.08 | 0.17 | 0.83 |
| MNM | 0.66 | 0.74 | 0.87 | 0.86 | 0.87 | 0.88 | 0.88 | 0.86 | 0.06 | 0.18 | 0.89 |
| Kmeans | 0.58 | 0.68 | 0.80 | 0.80 | 0.85 | 0.85 | 0.85 | 0.79 | 0.06 | 0.16 | 0.86 |
| SOM | 0.54 | 0.51 | 0.62 | 0.51 | 0.65 | 0.65 | 0.67 | 0.57 | 0.06 | 0.18 | 0.51 |

**Fig. S4 | Harmonization performance across sample sizes.** Heatmaps show the mean ARI for datasets with per-group sample sizes of: (A) 100, (B) 50, and (C) 27, under strong signal ($c = 1.5$) and weak artifacts ($d = 1.2$). Larger sample sizes yield only modest ARI gains and largely preserve the relative ranking of harmonization and clustering methods. Panel A is an independent rerun of the simulations shown in Fig. 4B-3 of the main text, yielding essentially identical values, and is included to facilitate visual comparison with panels B and C.

| (A) | Raw | TC | UQ | Med | TMM | DESeq | PoissonSeq | QN | RUVg | RUVr | RUVs |
|---|---|---|---|---|---|---|---|---|---|---|---|
| HC-E | 0.27 | 0.01 | 0.11 | 0.03 | 0.07 | 0.06 | 0.12 | 0.03 | 0.04 | 0.73 | 0.02 |
| HC-P | 0.05 | 0.03 | 0.06 | 0.02 | 0.05 | 0.03 | 0.05 | 0.04 | 0.03 | 0.19 | 0.00 |
| HC-S | 0.00 | 0.00 | 0.03 | 0.01 | 0.01 | 0.01 | 0.02 | 0.03 | 0.01 | 0.14 | 0.00 |
| PAM-E | 0.11 | 0.04 | 0.11 | 0.06 | 0.09 | 0.08 | 0.07 | 0.06 | 0.02 | 0.41 | 0.00 |
| PAM-P | 0.02 | 0.02 | 0.02 | 0.01 | 0.02 | 0.01 | 0.02 | 0.01 | 0.02 | 0.07 | 0.00 |
| PAM-S | 0.01 | 0.00 | 0.02 | 0.01 | 0.01 | 0.01 | 0.02 | 0.01 | 0.01 | 0.10 | 0.00 |
| MNM | 0.24 | 0.06 | 0.17 | 0.11 | 0.19 | 0.19 | 0.22 | 0.08 | 0.04 | 0.77 | 0.03 |
| Kmeans | 0.18 | 0.00 | 0.05 | 0.00 | 0.02 | 0.02 | 0.03 | 0.00 | 0.03 | 0.41 | 0.01 |
| SOM | 0.13 | 0.01 | 0.05 | 0.03 | 0.04 | 0.04 | 0.05 | 0.03 | 0.03 | 0.20 | 0.01 |

| (B) | Raw | TC | UQ | Med | TMM | DESeq | PoissonSeq | QN | RUVg | RUVr | RUVs |
|---|---|---|---|---|---|---|---|---|---|---|---|
| HC-E | 0.56 | 0.25 | 0.44 | 0.30 | 0.49 | 0.53 | 0.56 | 0.24 | 0.04 | 0.55 | 0.02 |
| HC-P | 0.51 | 0.44 | 0.45 | 0.42 | 0.42 | 0.47 | 0.45 | 0.34 | 0.07 | 0.50 | 0.04 |
| HC-S | 0.21 | 0.24 | 0.27 | 0.19 | 0.13 | 0.18 | 0.30 | 0.18 | 0.06 | 0.43 | 0.01 |
| PAM-E | 0.52 | 0.45 | 0.60 | 0.58 | 0.70 | 0.72 | 0.63 | 0.59 | 0.05 | 0.33 | 0.02 |
| PAM-P | 0.17 | 0.16 | 0.13 | 0.11 | 0.12 | 0.12 | 0.14 | 0.12 | 0.06 | 0.19 | 0.02 |
| PAM-S | 0.12 | 0.13 | 0.17 | 0.12 | 0.10 | 0.11 | 0.15 | 0.12 | 0.04 | 0.23 | 0.00 |
| MNM | 0.47 | 0.45 | 0.61 | 0.59 | 0.72 | 0.77 | 0.73 | 0.55 | 0.03 | 0.56 | 0.02 |
| Kmeans | 0.40 | 0.15 | 0.28 | 0.13 | 0.28 | 0.25 | 0.38 | 0.13 | 0.03 | 0.39 | 0.01 |
| SOM | 0.32 | 0.09 | 0.20 | 0.14 | 0.19 | 0.18 | 0.21 | 0.14 | 0.04 | 0.33 | 0.02 |

| (C) | Raw | TC | UQ | Med | TMM | DESeq | PoissonSeq | QN | RUVg | RUVr | RUVs |
|---|---|---|---|---|---|---|---|---|---|---|---|
| HC-E | 0.72 | 0.73 | 0.79 | 0.76 | 0.85 | 0.84 | 0.84 | 0.75 | 0.04 | 0.22 | 0.22 |
| HC-P | 0.88 | 0.84 | 0.92 | 0.90 | 0.92 | 0.94 | 0.92 | 0.91 | 0.07 | 0.22 | 0.53 |
| HC-S | 0.91 | 0.86 | 0.92 | 0.92 | 0.93 | 0.95 | 0.93 | 0.91 | 0.08 | 0.23 | 0.52 |
| PAM-E | 0.64 | 0.69 | 0.69 | 0.72 | 0.80 | 0.81 | 0.78 | 0.72 | 0.06 | 0.13 | 0.33 |
| PAM-P | 0.50 | 0.55 | 0.46 | 0.39 | 0.49 | 0.50 | 0.52 | 0.46 | 0.09 | 0.20 | 0.17 |
| PAM-S | 0.48 | 0.58 | 0.59 | 0.51 | 0.49 | 0.53 | 0.59 | 0.51 | 0.07 | 0.22 | 0.18 |
| MNM | 0.65 | 0.76 | 0.86 | 0.84 | 0.85 | 0.88 | 0.86 | 0.83 | 0.04 | 0.26 | 0.24 |
| Kmeans | 0.60 | 0.68 | 0.72 | 0.71 | 0.80 | 0.81 | 0.81 | 0.68 | 0.05 | 0.21 | 0.13 |
| SOM | 0.55 | 0.45 | 0.55 | 0.44 | 0.58 | 0.57 | 0.62 | 0.48 | 0.05 | 0.24 | 0.07 |

**Fig. S5 | Harmonization performance under cluster-size imbalance.** Heatmaps show the mean ARI as the MXF:UPS sample-size ratio departs from 1:1: (A) 1:9, (B) 1:4, and (C) 2:3, with the total sample size fixed at 200, under strong signal ($c = 1.5$) and weak artifacts ($d = 1.2$). Severe imbalance drives most ARIs toward zero, with a notable exception of RUVr combined with MNM.

**(A)**

| | Raw | TC | UQ | Med | TMM | DESeq | PoissonSeq | QN | RUVg | RUVr | RUVs |
|---|---|---|---|---|---|---|---|---|---|---|---|
| HC-E | 0.53 | 0.61 | 0.66 | 0.66 | 0.67 | 0.67 | 0.67 | 0.66 | 0.08 | 0.15 | 0.63 |
| HC-P | 0.55 | 0.55 | 0.54 | 0.54 | 0.54 | 0.54 | 0.54 | 0.54 | 0.09 | 0.13 | 0.52 |
| HC-S | 0.60 | 0.56 | 0.55 | 0.54 | 0.54 | 0.54 | 0.55 | 0.54 | 0.10 | 0.15 | 0.54 |
| PAM-E | 0.45 | 0.54 | 0.48 | 0.50 | 0.56 | 0.55 | 0.55 | 0.52 | 0.07 | 0.08 | 0.53 |
| PAM-P | 0.35 | 0.46 | 0.44 | 0.43 | 0.45 | 0.46 | 0.46 | 0.45 | 0.10 | 0.12 | 0.46 |
| PAM-S | 0.38 | 0.46 | 0.46 | 0.46 | 0.48 | 0.48 | 0.46 | 0.46 | 0.09 | 0.12 | 0.48 |
| MNM | 0.57 | 0.58 | 0.68 | 0.70 | 0.74 | 0.71 | 0.68 | 0.69 | 0.09 | 0.18 | 0.69 |
| Kmeans | 0.46 | 0.59 | 0.61 | 0.63 | 0.63 | 0.62 | 0.62 | 0.62 | 0.10 | 0.13 | 0.58 |
| SOM | 0.30 | 0.38 | 0.37 | 0.31 | 0.39 | 0.39 | 0.41 | 0.35 | 0.03 | 0.11 | 0.23 |

**(B)**

| | Raw | TC | UQ | Med | TMM | DESeq | PoissonSeq | QN | RUVg | RUVr | RUVs |
|---|---|---|---|---|---|---|---|---|---|---|---|
| HC-E | 0.43 | 0.54 | 0.55 | 0.57 | 0.57 | 0.56 | 0.56 | 0.55 | 0.08 | 0.13 | 0.52 |
| HC-P | 0.36 | 0.37 | 0.37 | 0.37 | 0.37 | 0.37 | 0.37 | 0.37 | 0.07 | 0.10 | 0.36 |
| HC-S | 0.38 | 0.37 | 0.37 | 0.37 | 0.38 | 0.38 | 0.38 | 0.38 | 0.08 | 0.11 | 0.37 |
| PAM-E | 0.36 | 0.39 | 0.36 | 0.39 | 0.45 | 0.43 | 0.40 | 0.41 | 0.07 | 0.07 | 0.44 |
| PAM-P | 0.28 | 0.32 | 0.31 | 0.31 | 0.33 | 0.33 | 0.33 | 0.32 | 0.08 | 0.09 | 0.32 |
| PAM-S | 0.31 | 0.32 | 0.32 | 0.32 | 0.34 | 0.34 | 0.33 | 0.32 | 0.08 | 0.09 | 0.34 |
| MNM | 0.49 | 0.57 | 0.61 | 0.63 | 0.65 | 0.63 | 0.62 | 0.62 | 0.09 | 0.14 | 0.61 |
| Kmeans | 0.39 | 0.51 | 0.50 | 0.53 | 0.53 | 0.51 | 0.50 | 0.50 | 0.09 | 0.12 | 0.48 |
| SOM | 0.22 | 0.36 | 0.31 | 0.29 | 0.33 | 0.33 | 0.34 | 0.31 | 0.03 | 0.08 | 0.20 |

**(C)**

| | Raw | TC | UQ | Med | TMM | DESeq | PoissonSeq | QN | RUVg | RUVr | RUVs |
|---|---|---|---|---|---|---|---|---|---|---|---|
| HC-E | 0.36 | 0.46 | 0.45 | 0.47 | 0.47 | 0.44 | 0.43 | 0.45 | 0.07 | 0.11 | 0.44 |
| HC-P | 0.27 | 0.28 | 0.28 | 0.27 | 0.28 | 0.28 | 0.28 | 0.28 | 0.06 | 0.08 | 0.27 |
| HC-S | 0.29 | 0.28 | 0.28 | 0.28 | 0.29 | 0.29 | 0.28 | 0.28 | 0.07 | 0.09 | 0.28 |
| PAM-E | 0.30 | 0.33 | 0.32 | 0.33 | 0.39 | 0.36 | 0.34 | 0.35 | 0.06 | 0.06 | 0.38 |
| PAM-P | 0.23 | 0.25 | 0.25 | 0.24 | 0.25 | 0.26 | 0.25 | 0.25 | 0.06 | 0.07 | 0.25 |
| PAM-S | 0.24 | 0.25 | 0.24 | 0.25 | 0.26 | 0.26 | 0.26 | 0.25 | 0.06 | 0.07 | 0.26 |
| MNM | 0.43 | 0.54 | 0.54 | 0.56 | 0.58 | 0.56 | 0.54 | 0.56 | 0.07 | 0.12 | 0.54 |
| Kmeans | 0.33 | 0.41 | 0.39 | 0.41 | 0.42 | 0.39 | 0.37 | 0.40 | 0.08 | 0.11 | 0.41 |
| SOM | 0.18 | 0.30 | 0.27 | 0.28 | 0.28 | 0.28 | 0.28 | 0.27 | 0.03 | 0.06 | 0.19 |

**Fig. S6 | Harmonization performance under cluster-number misspecification.** Heatmaps show the mean ARI when the number of clusters is over-specified: (A) K = 4; (B) K = 6; (C) K = 8, under strong signal ($c$ = 1.5) and weak artifacts ($d$ = 1.2); the true number of clusters is 2 for a set of 100 MXF samples and 100 UPS samples. Over-partitioning consistently degrades ARI, most severely for correlation-based HC-P and HC-S, more moderately for HC-E, and least for the model-based MNM.

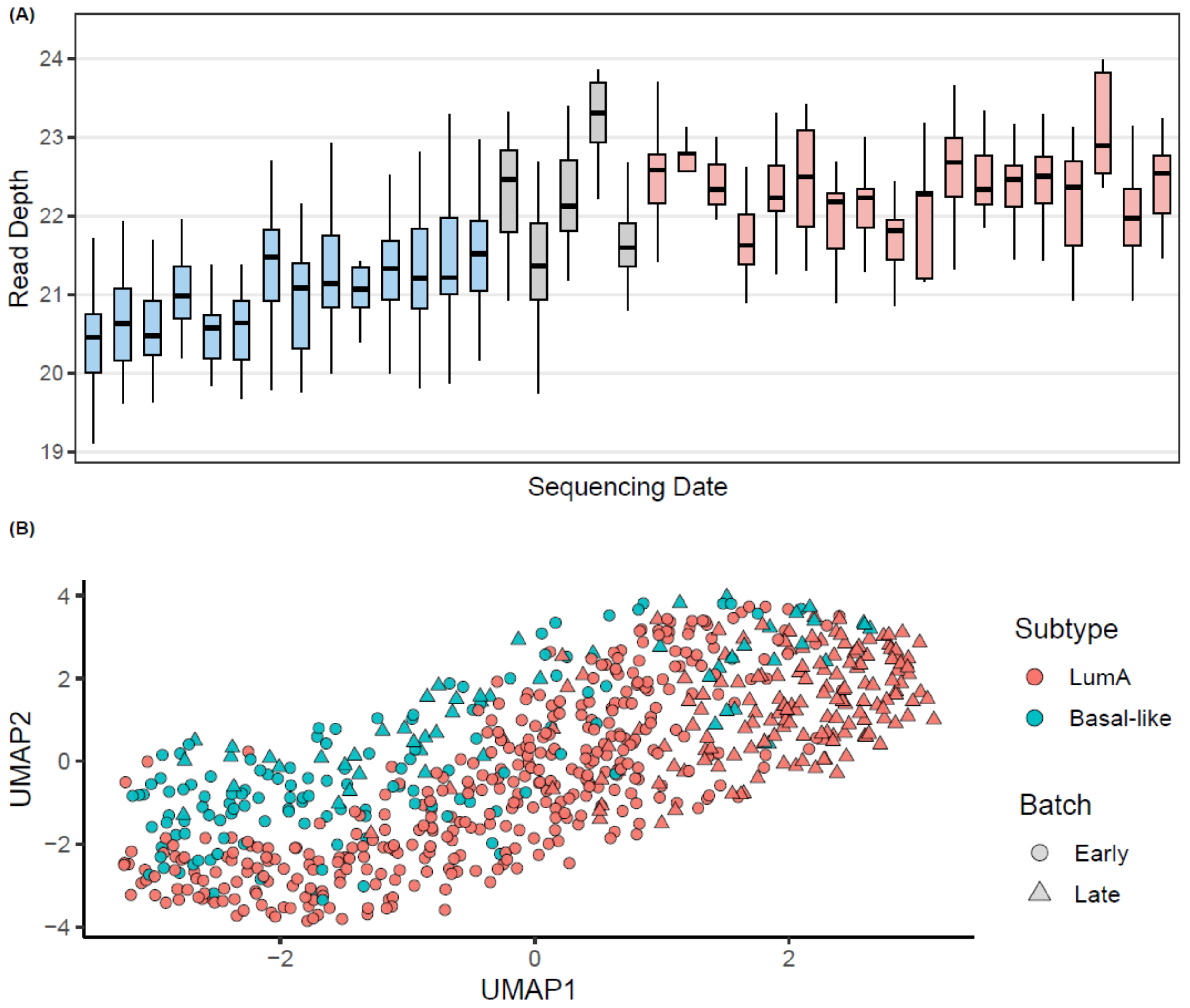


**Fig. S7 | Definition of data collection batches and their association with tumor subtype in the TCGA breast cancer study.** (A) Boxplots of sequencing depth (i.e., total read count per sample) over data collection date, revealing an Early batch (blue) and a Late batch (red), separated by a transition period (grey) that was excluded from cohort reconstruction. (B) UMAP of log2-transformed sequencing depth, with each point representing a sample. Colors indicate breast-cancer subtypes (Luminal A vs. Basal-like) and shapes denote sequencing data batch.

(A)

| | LumA | Basal |
|---|---|---|
| Early | 0 | 0 |
| Late | 50 | 50 |

(B)

| | LumA | Basal |
|---|---|---|
| Early | 25 | 25 |
| Late | 25 | 25 |

(C)

| | LumA | Basal |
|---|---|---|
| Early | 0 | 50 |
| Late | 50 | 0 |

(D)

| | LumA | Basal |
|---|---|---|
| Early | 50 | 0 |
| Late | 0 | 50 |

**Table S1 | Sample-size allocations by subtype and batch for the reconstructed cohorts from the TCGA breast cancer study under four artifact scenarios.** (A) Scenario I – weak artifact, with all samples drawn from the late batch; (B) Scenario II – strong, balanced artifacts, with equal numbers of early- and late-batch samples for each subtype; (C) Scenario III – strong, completely confounded artifacts, with all Luminal A samples from the early batch and all Basal samples from the late batch; and (D) Scenario IV – same as Scenario III but with batch assignment switched between subtypes.
